# Extensive Novel Genomic Variations in Mutant European Pear Individuals Revealed by Mapping to a Pangenome Reference

**DOI:** 10.64898/2026.03.02.709077

**Authors:** June Labbancz, Nathan Tarlyn, Kate Evans, Amit Dhingra

## Abstract

European pear (*Pyrus communis*) is the most widely cultivated *Pyrus* species outside of Asia; however, production has been declining in Europe, Oceania, and North America. Most European pear cultivars are over 100 years old and face pressure from disease, a changing climate, and challenging postharvest characteristics, while demand for increased production efficiency rises. To generate germplasm with desirable characteristics to address these concerns, a mutation breeding approach was chosen, with crosses made between four economically significant cultivars (‘Bartlett’, ‘d’Anjou’, ‘Abbe Fetel’, and ‘Comice’) using gamma-irradiated pollen. This resulted in 49 viable offspring, of which 37 have survived at least 10 years. Nanopore whole-genome sequencing was used to test the success of this approach and screen for variants of interest. Sequence reads were mapped to both a lightweight, purpose-built pangenome derived from assemblies of parental haplotypes and a linear reference genome, enabling the high-quality discovery of variants of all sizes, ranging from single-base substitutions to megabase-scale deletions, with the overwhelming majority being small variants. The overall rate of mutation was 153 novel small variants and 0.228 novel structural variants per Gray of absorbed gamma radiation. Alternate ploidy levels were detected in four lines, which included three triploids and one tetraploid. While the resulting individuals appear incapable of floral development, they may be of utility as rootstock cultivars and a valuable genetic resource for understanding the underlying basis of structural traits.

## Introduction

Despite the economic importance of European pear (*Pyrus communis*) cultivation in North America and Europe, the majority of cultivation uses scion cultivars that are over 100 years old. In North America, ‘Bartlett’ and ‘d’Anjou’ dominate production (Northwest Horticultural Council, 2025), while in Europe, ‘Conference’ has become increasingly popular, with ‘Bartlett’ and ‘Abate Fetel’ also contributing to most of the production. While some semi-dwarfing rootstocks have become available, notably the ‘Pyrodwarf’ and ‘Old Home’ × ‘Farmingdale’ lines, most pears are still grown using non-dwarfing rootstocks or non-*Pyrus* rootstocks, with options being limited compared to those in *Malus* (Brewer and Palmer, 2011). Using quince (*Cydonia oblonga*) as a rootstock for European pear cultivation is the most common means of inducing precocity and controlling scion vigor, enabling high-density plantingsClick or tap here to enter text., however quince rootstocks can suffer from graft incompatibility, iron chlorosis, and poor tolerance of hot climates (Musacchi et al., 2021). Most European pear cultivars are vulnerable to many pests, diseases, and environmental pressures. Fire Blight (*Erwinia amylovora*), for example, has vastly reduced pear production in the Eastern United States, and pear production in Europe has shifted northwards, in response to climate-related crop losses in Southern growing regions (Gottschalk et al., 2024; Musacchi, 2024). Difficulties are not limited to production; the chilling requirements for pear ripening, the fragility of ripe pears, and the incompatibility of pears with 1-methylcyclopropene (1-MCP) ethylene inhibitor treatment make delivering a consistent consumer eating experience difficult (Hewitt et al., 2020; Tedeschi et al., 2023). Sustainable pear production, which also meets consumer expectations for ripe, flavorful fruit year-round, is increasingly out of the grasp of the pear industry (Musacchi, 2024); consequently, production of European pear has decreased over the past 20 years in Europe and the Americas by approximately 30% and 10%, respectively (Food and Agriculture Organization of the United Nations, 2024).

Other *Pyrus* species have desirable traits which are lacking in existing European pear germplasm, however, incorporating these species into breeding programs is impractical for scion cultivars, as they are likely to negatively impact the highly desirable traits of European pear fruit, such as its large size, melting texture, or balanced sugar to acid content (Evrenosoğlu and Mertoğlu, 2018; Zhang et al., 2025). This is especially problematic in the context of the high heterozygosity of *Pyrus* individuals. Cultivars resulting from the hybridization of European and Asian pears (*P. pyrifolia, P. ussuriensis,* and *P. bretschneideri*) are numerous but have not been successful in the market. Development of new rootstock cultivars offers greater potential for the incorporation of other *Pyrus* species, yet vigor-controlling precocious rootstock options remain sparse for European pear (Brewer and Palmer, 2011; Guzman and Dhingra, 2019).

While the European pear is generally propagated clonally, mutagenesis of somatic tissues can lead to chimerism, which can pose challenges in the development of new cultivars (Jankowicz-Cieslak and Till, 2017), and genome editing (Medvedieva and Blume, 2018; Vora et al., 2023), application is limited in the near term by lack of knowledge of the molecular basis of most pear traits For these reasons, mutation breeding is particularly suited to the development of novel European pear genetic resources; mutagenesis of reproductive tissues such as pollen prior to breeding may be an expedient approach to generate observable phenotypes in the F1 generation.

Irradiation typically results not only in single-nucleotide polymorphisms and other small variants, which are easily captured by short-read sequencing, but also in a small number of larger structural variants that may span thousands of base pairs. The detection of the latter type of genomic changes requires long-read sequencing for adequate detection (Stephens et al., 2018; Kitamura et al., 2022; Youk et al., 2024). The exact frequency and types of mutations may vary considerably depending on experimental design, with tissue type, tissue maturity, radiation dose, duration of radiation exposure, and radiation source all influencing the resulting mutation pattern (Kazama et al., 2017; Li et al., 2019a; Hase et al., 2020; Choi et al., 2021). Gamma irradiation tends to induce smaller variants on average than ion beam irradiation, for example (Hase et al., 2020).

While alignment of sequencing reads to single linear reference genomes has enabled much of modern genomics, such studies are likely to suffer from reference bias, a phenomenon where genetic variants may only be discoverable when they are sufficiently similar to the reference sequence, and from poor sensitivity in variant-rich areas (Miga and Wang, 2020; Du et al., 2025; Hu et al., 2025). This is particularly pronounced in situations where common structural variants are omitted from the reference genome. While long read sequencing ameliorates some of these issues, increasing the likelihood of sequence reads spanning entire variants, helping especially in structural variant detection, alignment to pangenome sequence graphs composed of many individuals improves variant detection (Kosugi and Terao, 2024; Mahmoud et al., 2024; Schloissnig et al., 2025). Recent advances in sequencing technology and the development of pangenomic resources for a broader range of species and genera enable the accurate genetic evaluation of individuals produced through mutagenesis.

While pangenome graphs and existing variant calling tools, which have graph compatibility, excel at identifying common variants within a population, especially relative to linear references, the identification of novel variants is considerably more difficult (Du et al., 2025). This poses a significant challenge for the analysis of mutation-bred individuals in particular, that are likely to have thousands of novel variants. Additionally, while all existing graph generation pipelines are useful for detecting structural variation, not all pipelines consider smaller variants, which may be more prone to reference bias. Minigraph, the most highly cited pangenome graph generation tool for eukaryotic genomes, omits variants smaller than 50bp by default (Li et al., 2020). The Minigraph-Cactus and Pangenome Graph Builder (PGGB) pipelines are considered lossless and aim to encode all variation, but may become unwieldy or fail entirely, especially for large numbers of haplotypes and in lineages with high degrees of variation (Andreace et al., 2023; Heumos et al., 2024; Hickey et al., 2024). Selecting a limited set of individuals may help overcome some of these scalability concerns; in a breeding context where a small number of parents with desirable traits have been preselected, this approach may be preferred. While effective tools for genotyping known structural variants in a population have been developed (Sirén et al., 2021), there is a paucity of tools that enable the scalable identification of novel structural variants using a pangenome reference. However, tools for the identification of novel small variants, such as the vg toolkit, have been developed (Hickey et al., 2020).

With the goal of producing novel variation within elite European pear cultivars, crosses were made between four widely cultivated commercial pear cultivars (Bartlett’, ‘d’Anjou’, ‘Comice’, and ‘Abbe Fetel’) using gamma-irradiated pollen in each cross (Table 1). Putative mutant seeds resulting from these crosses were germinated and propagated once at an appropriate size. To validate successful mutagenesis, DNA purified from leaf tissue representing each surviving mutation-bred line was sequenced using Nanopore whole-genome sequencing. A small pangenome graph consisting only of 8 haplotypes, 2 per parental accession, was constructed to facilitate analysis. Long sequencing reads were then mapped to a pangenome graph for *Pyrus* (Labbancz et al., Unpublished), enabling the characterization of resulting small (<50bp) novel genetic variants. Mapping to a linear reference genome was used for the detection of structural variants (>50bp) and copy number variants.

**Table 1.** Recorded genetic background of all lines sequenced. The first name indicates the mother (emasculated flower), and the second name, followed by an “(i)”, indicates the father (irradiated pollen). The genetic background of Sample 31 was lost due to degradation of the identifier tag, indicating the parents and cross.

| Sample | Cross | Identifier |
| --- | --- | --- |
| 1 | 'Bartlett' x 'd'Anjou'(i) | 13-1 |
| 2 | 'Bartlett' x 'd'Anjou'(i) | 13-3 |
| 3 | 'Bartlett' x 'd'Anjou'(i) | 13-6 |
| 4 | 'Bartlett' x 'Bartlett'(i) | 13-1 |
| 5 | 'Bartlett' x 'Bartlett'(i) | 13-2 |
| 6 | 'Bartlett' x 'Bartlett'(i) | 13-3 |
| 7 | 'Bartlett' x 'Bartlett'(i) | 13-5 |
| 8 | 'Bartlett' x 'Bartlett'(i) | 13-6 |
| 9 | 'Bartlett' x 'Bartlett'(i) | 13-7 |
| 10 | 'Bartlett' x 'Comice'(i) | 13-1 |
| 11 | 'Bartlett' x 'Comice'(i) | 13-3 |
| 12 | 'Bartlett' x 'Abbe Fetel'(i) | 13-1 |
| 13 | 'Bartlett' x 'Abbe Fetel'(i) | 13-10 |
| 14 | 'Bartlett' x 'Abbe Fetel'(i) | 13-11 |
| 15 | 'Bartlett' x 'Abbe Fetel'(i) | 13-12 |
| 16 | 'Bartlett' x 'Abbe Fetel'(i) | 13-14 |
| 17 | 'Bartlett' x 'Abbe Fetel'(i) | 13-15 |
| 18 | 'Bartlett' x 'Abbe Fetel'(i) | 13-2 |
| 19 | 'Bartlett' x 'Abbe Fetel'(i) | 13-3 |
| 20 | 'Bartlett' x 'Abbe Fetel'(i) | 13-5 |
| 21 | 'Bartlett' x 'Abbe Fetel'(i) | 13-9 |
| 22 | 'Comice' x 'd'Anjou'(i) | 13-1 |
| 23 | 'Comice' x 'Comice'(i) | 13-1 |
| 24 | 'Comice' x 'Comice'(i) | 13-2 |
| 25 | 'Comice' x 'Comice'(i) | 13-4 |
| 26 | 'Comice' x 'Comice'(i) | 13-5 |
| 27 | 'Comice' x 'Comice'(i) | 13-8 |
| 28 | 'Abbe Fetel' x 'Abbe Fetel'(i) | 13-1 |
| 29 | 'Abbe Fetel' x 'Abbe Fetel'(i) | 13-2 |
| 30 | 'Abbe Fetel' x 'Abbe Fetel'(i) | 13-3 |
| 31 | Illegible |  |
| 32 | 'd'Anjou' x 'd'Anjou'(i) | 2017 |
| 33 | 'Bartlett' x 'Abbe Fetel'(i) | 13-6 |
| 34 | 'Bartlett' x 'Abbe Fetel'(i) | 13-7 |
| 35 | 'Bartlett' x 'd'Anjou'(i) | 13-8 |
| 36 | 'Bartlett' x 'd'Anjou'(i) | 13-9 |
| 37 | 'Bartlett' x 'd'Anjou'(i) | 13-12 |

## Methods

### Mutagenesis

Pollen was collected from the four parent cultivars ‘Bartlett’, ‘d’Anjou’, ‘Comice’, and ‘Abate Fetel’ in Spring 2013. Gamma irradiation was conducted at the Washington State University Nuclear Science Facility using a Cobalt-60 source. Pollen received a total dose of 189 kilorads at a dose of 0.45 kilorads/minute over a period of 7 hours. Flowers pollenated with irradiated pollen were emasculated prior to the application of irradiated pollen, and pollenated flowers were bagged to prevent entry of non-irradiated pollen. Fruit from the crosses were marked and harvested at maturity. The resulting seeds were stratified in petri dishes lined with damp paper towels for 90 days at 4°C. Germinated seedlings were transferred to Sunshine #4 potting mix (Sun Gro Horticulture, Agawam, MA, USA) in 50-cell plug trays. One sample (#32) was produced by the same method in 2017.

### Seedling Growth Conditions

Post-germination, plants were transferred to a greenhouse with supplemental lighting to maintain a 12-hour or greater photoperiod through the winter. Temperature was maintained at 22-26°C. Once seedlings grew to a height of 10-15cm, they were transferred to ∼160mL tree pots. To meet European pear chilling requirements, 1000 hours of chilling at 0-4°C was provided in a climate-controlled room annually. Once saplings had achieved a trunk diameter of 6-7mm, they were moved to 2-gallon pots (5-gallon for sample #32).

Trees were propagated *in vitro*. Nodal segments were sterilized by rinsing with water for at least one hour, followed by a 1-minute wash in 70% ethanol, then a 5-minute wash in 0.1% mercury chloride+0.01% Silwet L-77. Nodal segments were placed in a sterile media produced by the mixture of 1X Quoirin & Lepoivre Media basal salts, 250mg/L MES [2-(N-morpholino)-ethanesulfonic acid], 100mg/L myo-inositol, 16g/L glucose, 1mg/L 6-Benzylaminopurine, 0.1mg/L indole-3-butyric acid (IBA), adjusted to a pH of 5.8. As shoots developed, multiplication of explants was achieved by dividing nodal segments and placing into fresh sterile media. Explants were prepared for rooting by dipping the bottom 1cm of stem in 10mg/mL IBA in DMSO, then placed on a phytohormone-free version of the sterile media for propagation.

Once roots formed, explants were removed from the sterile media and transferred to damp potting soil with a plastic humidity dome. After at least one week of establishment, the humidity dome was removed.

Of these clonally propagated individuals, some were maintained in 2-gallon pots, while others were planted at the Washington State University Horticulture Center, Pullman, WA. In 2022, samples #1-31 were transferred from Pullman, WA, to the Horticulture Teaching, Research, and Extension Center in Somerville, TX, in 2-gallon pots. In 2024, sample #32 was transferred from Pullman, WA, to a greenhouse in College Station, TX. In 2024, scion cuttings were taken from samples #33-37 at the Washington State University Horticulture Center, Pullman, WA, and cleft grafted to *P. betulafolia* rootstocks in College Station, TX. At the time of tissue collection, all trees were maintained at the Horticulture Teaching, Research, and Extension Center in Somerville, TX, in 3-gallon pots in LP15 potting mix (Sun Gro Horticulture, Agawam, MA, USA). In the absence of adequate chilling hours for European pears at this location, at least 6 weeks of chilling at 0-4°C were provided in a climate-controlled room annually.

### Sample Processing

Leaf samples were collected from tender tissue within one month of bud break from the putative mutants in Spring 2024 and Spring 2025. A total of 3-5 g of leaf material was collected per line, immediately placed in a 50mL polypropylene tube, and immersed in liquid nitrogen. Leaf material was cryomilled using a Spex Sample Prep Freezer/Mill 6875 (Metuchen, NJ, USA) for 2 minutes at a rate of 15cpm. Milled samples were stored at –80 °C until DNA isolation.

Following crude DNA isolation using a modified CTAB protocol, the DNA was precipitated with an equal volume of isopropanol and incubated overnight at 4 °C. The precipitated DNA was then washed twice with 75% ethanol, with centrifugation at 12,000g for 5 minutes at 4 °C between washes. DNA was resuspended in 5mL 1.56g/mL CsCl_2_ solution at 42°C overnight to ensure complete resuspension. A total of 5µg Ethidium Bromide or 1 µL 10,000X SYBR Gold was added to the sample, loaded into Eppendorf 5PP seal tubes, and brought within 100mg of one another using 1.56g/mL CsCl solution before thermal sealing and centrifugation.

Resuspended DNA was ultracentrifuged for 6 hours at 70,000rpm in a Himac P100VT rotor (Himac, Hitachinaka, Japan). DNA was extracted from the 5PP seal tubes using an 18-gauge needle. The extracted samples were diluted with three volumes of sterile MilliQ (Millipore Sigma, Billerica, MA, USA) water and precipitated with an equal volume of isopropanol. The DNA pellet was washed twice with 75% ethanol, and the samples were resuspended in 1X TE (10mM Tris, 1mM EDTA, pH 8.0) overnight at 42°C. Subsequently, the Oxford Nanopore Short Fragment Exclusion Kit (Oxford Nanopore, Oxford, United Kingdom) was used to remove short DNA fragments and select for longer DNA fragments, using the prescribed protocol, with the sole modification of extending the main centrifugation step to 45 minutes from 30 minutes. The purity of isolated DNA was estimated by using a NanoDrop 8000 UV spectrophotometer, and quantity was determined using a QuBit 4 in conjunction with the 1X dsDNA BR working buffer.

### DNA Sequencing

Size-selected total DNA was prepared into multiple sequencing libraries using the Oxford Nanopore Native Barcoding 24 Kit (Oxford Nanopore, Oxford, United Kingdom) following the manufacturer’s protocol. Sequencing was conducted using R10.4.1 Flow Cells in conjunction with a Promethion P2 Solo device (Oxford Nanopore, Oxford, United Kingdom). Raw output was converted to .fastq reads using the “basecall” function of the Dorado software package, with a quality filter of Q10.

### Bioinformatic Analysis

A subset of *Pyrus* pangenome accessions (Labbancz et al. Unpublished) corresponding to the parents used in the breeding crosses in this experiment (*P. communis* cv. cultivars ‘Bartlett’, ‘d’Anjou’, ‘Comice’, and ‘Abate Fetel’) were used to generate an experiment-specific pangenome reference. A total of 8 haplotypes corresponding to 4 accessions were processed in the Minigraph-Cactus version 2.9.9 pangenome pipeline using default parameters (Hickey et al., 2024). Haplotype 1 of *Pyrus communis* cv. ‘Bartlett’ from this set was used as the reference sequence for all analyses to ensure shared reference coordinates.

Reads shorter than 10,000bp were filtered out using SeqKit v2.9.0 to improve the performance of alignment to the graph (Shen et al., 2024). Filtered reads were aligned to the pangenome graph using GraphAligner version 1.0.20, and variant calling was performed using the vg toolkit; vg giraffe aligned to the graph, vg augment added novel variants to the graph, vg pack was used to prepare the alignments for variant calling, and vg call was used to make variant calls and produce a .vcf file (Garrison et al., 2018; Hickey et al., 2020; Rautiainen and Marschall, 2020; Liao et al., 2023). Default parameters were used for all steps in the vg pipeline. Only variants >50BP and with a quality >50 were retained, and multiallelic calls were split. Bcftools version 1.22 isec was used to determine which variants were novel in each sample, not present in the vcf decomposition of the original parental haplotype graph (Danecek et al., 2021).

To detect large structural variants, a linear-based pipeline was used, using the same reference as the pangenome for consistency. Minimap2 was used with the flag -x map-ont to produce SAM alignment files, and Samtools was used to convert these files into sorted, indexed BAM files (Li, 2018; Danecek et al., 2021). Sniffles2 was used to call structural variants from the BAM alignment files, with an elevated mapq threshold of 30. As copy number variation was handled by another tool, deletions and duplications >200kb were filtered from the resulting vcf file.

SURVIVOR was used to merge structural variant VCF files, before the calling of variants unique to one sample from the merged vcf using bcftools view (Jeffares et al., 2017). These variants unique to only a single sample were deemed novel.

Copy number variants were assessed using CNVpytor version 1.3.1 (Suvakov et al., 2021). Alignments to the linear reference genome were filtered of all low-quality, secondary, and supplementary alignments using Samtools view with the flags (-F 0×904 -q 30). The reference genome mask and GC content per bin was packed into a CNVpytor-specific configuration file to avoid spurious variant calls in repetitive regions and regions with abnormal GC-content. Filtered bam files were processed by CNVpytor at two bin scales, 25kb and 100kb. Deletions present at both scales, with t-test p values < 0.001 and dosage ratios of 0.4-0.6, were retained. Duplications present at both scales, with t-test p values < 0.001 and dosage ratios of 1.4-1.6, were retained. The 25kb scale was used to determine the copy number variation length.

Ploidy analysis was performed by two means. First, Dorado corrected reads underwent k-mer counting with Jellyfish (at k=21), followed by k-mer histogram analysis with GenomeScope version 2.0 (Marçais and Kingsford, 2011; Ranallo-Benavidez et al., 2020). Second, BAM files generated during alignment for structural variant calling were used as inputs for nQuire to estimate ploidy using analysis of variant sites (Weiß et al., 2018).

Representative samples (#6, #25, and #37) were assembled using Hifiasm and mapped to the reference genome as a means of evaluating and validating variants. Hifiasm was run using the ‘--ont’ flag, and minimap2 was run using the ‘-x asm20’ flag. Samtools was used to convert outputs to BAM format, sort, and index. IGV was used to visualize all alignments (Robinson et al., 2023). BAM files used for structural variant detection were also used for evaluation and validation in conjunction with IGV. One gene at the *S*-locus (predicted gene chr17-g2006 – an *S*-locus F-box brother gene) was used to assess the *S*-haplotype, which is responsible for self-incompatibility (Claessen et al., 2019), to assess parentage and the potential breakdown of self-incompatibility by novel genetic variants. Parental assemblies and sequencing reads from each sample were aligned to the reference assembly using minimap2 (-x asm20 for parental accessions, -x map-ont for reads), then visualized in IGV for manual variant review (Robinson et al., 2017). First, SNPs for the parental *S*-allele were determined, and then the sample data were compared to these identified SNPs. In order to determine correlation between gene content and variant density, Pearson’s correlation was calculated for counts of small, structural, and copy number variants per 50kb window of the genome against the proportion of genic sequence for each 50kb window of the genome.

## Results and Discussion

### Mutation Pattern

Extensive novel mutations of varying sizes were detected in all samples sequenced in this experiment, including small variants, larger structural variants, and megabase-scale deletions (Figure 1). Small (< 50 bp) variants were detected in large numbers on all chromosomes in all samples in this study, with a median of 190,131 novel small variations not found in the original pangenome graph in each sample (Figure 2). Base substitutions are the most common small variant in most samples, while deletions vastly outnumber insertions in all cases (Figure 2, Figure 3a). Gamma irradiation tends to produce a profile of numerous small variants, with a bias towards small deletions and base substitutions (Hase et al., 2020; Hirao et al., 2022). While small variants were widely distributed across the genome on a large scale, at a finer scale, it was apparent that some regions had a high degree of small variants, while others were relatively free of variants (Figure 4).

**Figure 1.**
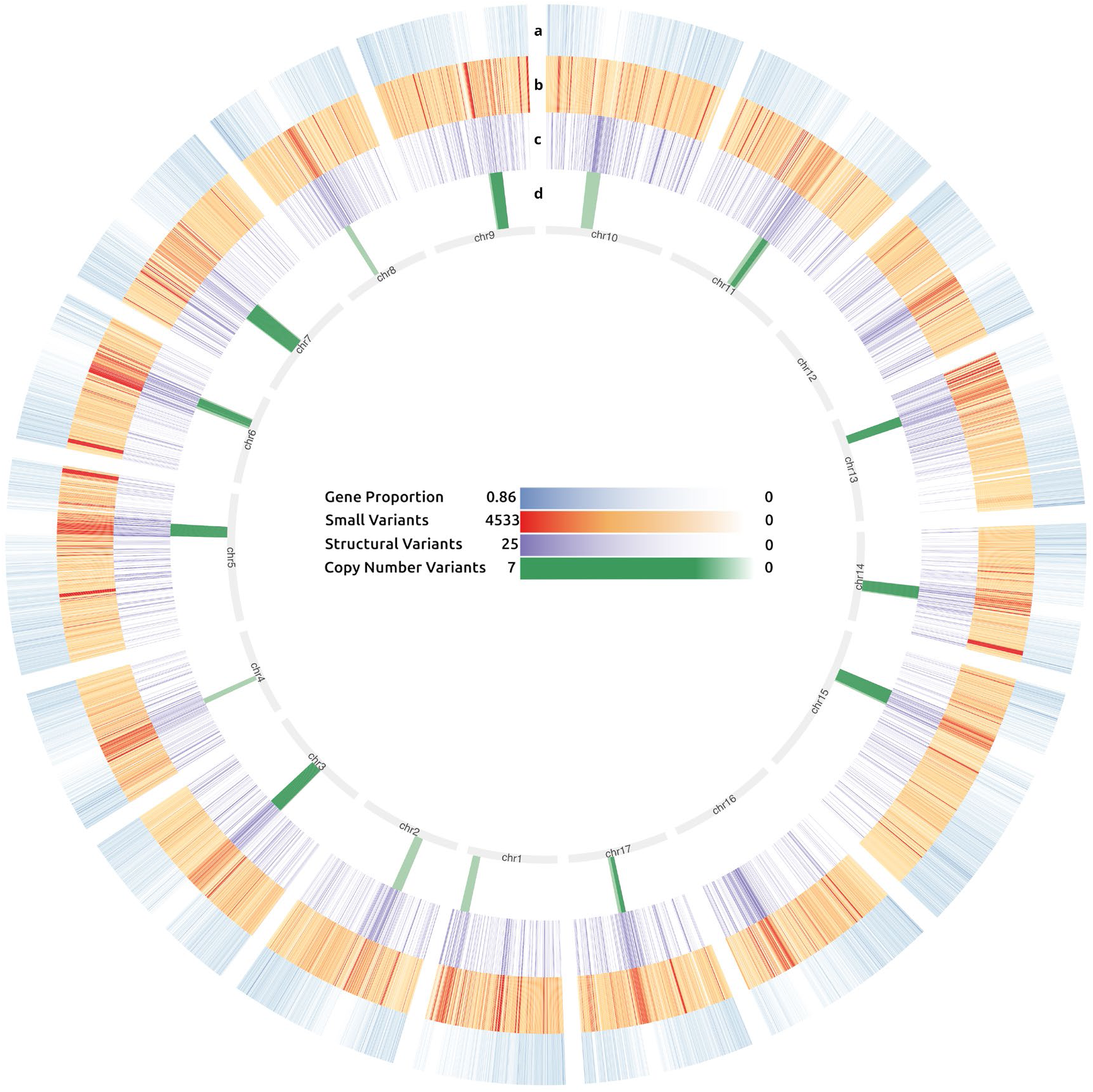
Circos plot of genic sequence proportion and variant density relative to the ‘Bartlett’ reference genome. Tracks correspond to a) relative gene density expressed as a proportion of the bin covered by a gene, b) relative small (<=50bp) variant frequency, c) relative structural variant (>50bp) frequency, and d) relative CNV-scale (>200kb) deletion frequency at 50kb bin resolution, with darker shading indicating higher density.

**Figure 2.**
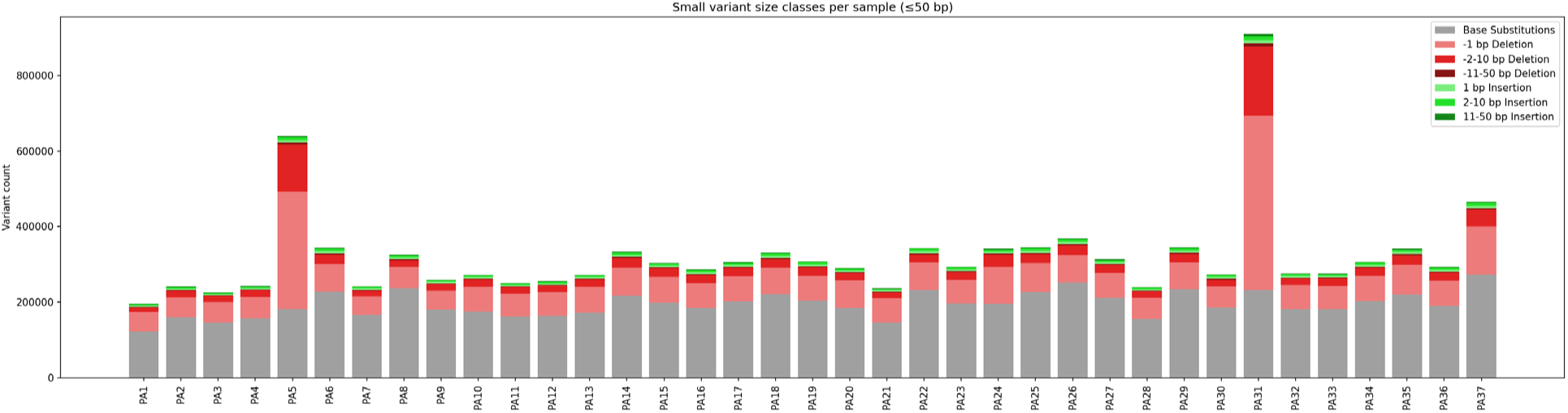
Chart displaying the number of small (<=50bp) variants per sample, divided by type. Base substitutions were predominantly represented in each sample. Most samples show comparable levels of variation.

**Figure 3:**
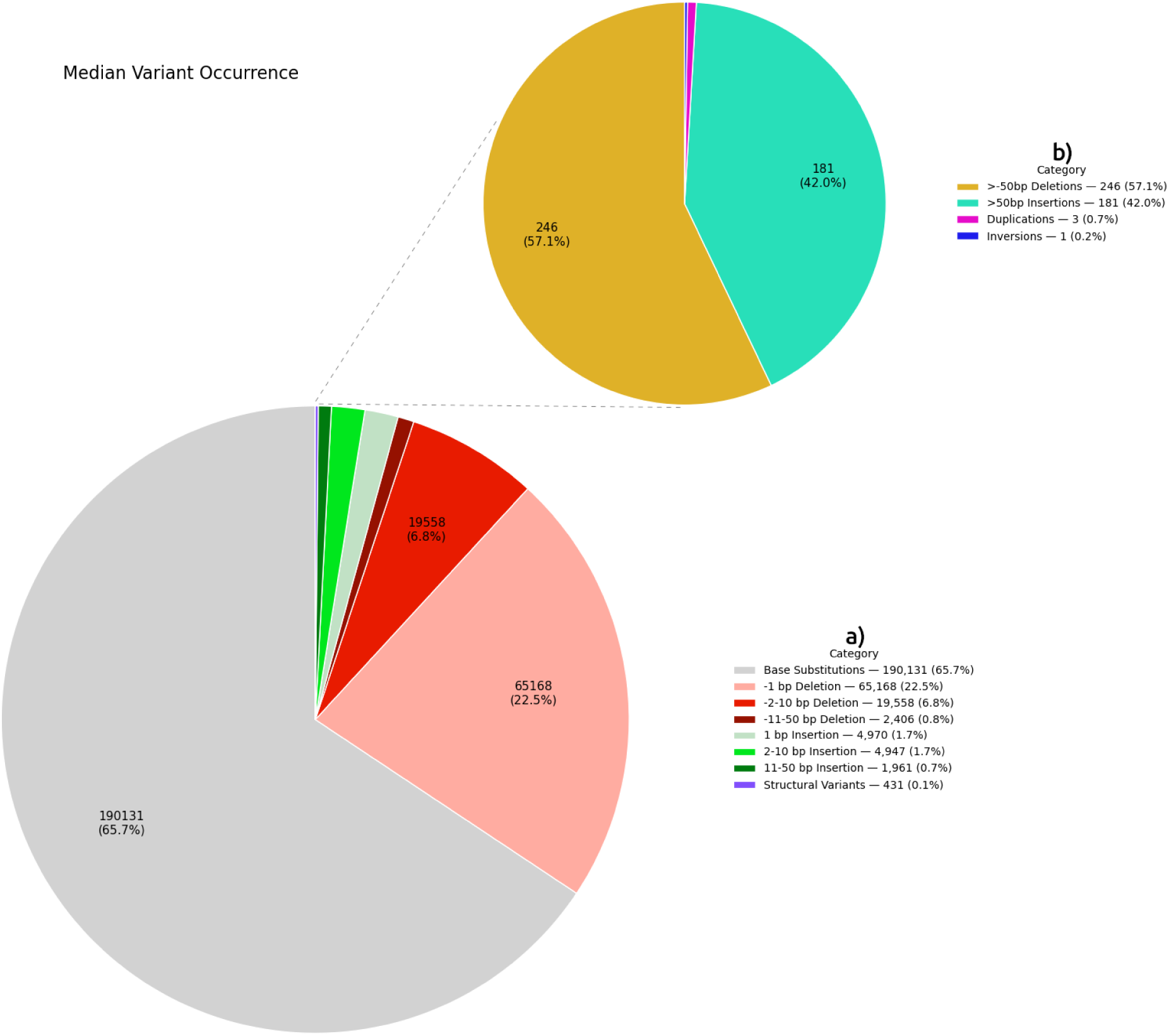
Plot of variant types at all scales. Median occurrence of a) small (<=50bp) variants divided by type, and b) structural variants (>50bp) by type. Samples #5 and #31 have been excluded, as these samples have at least one unknown parent which were not part of the original experimental design.

**Figure 4.**
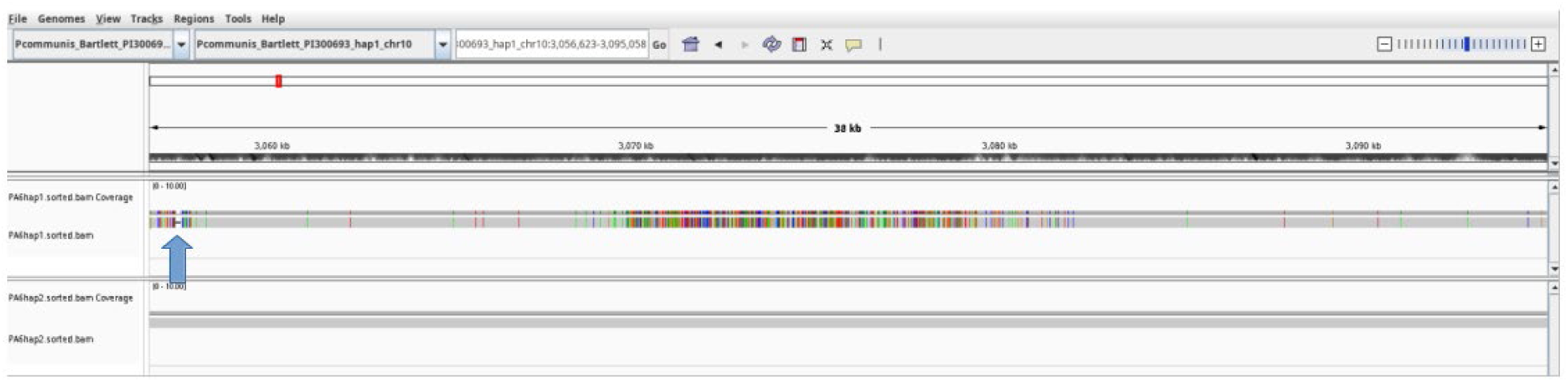
An IGV view displaying heterogeneity in the distribution of small variants within a sample. One deletion is represented by a black bar (indicated by a blue arrow), while base substitutions are represented by color. Some areas accumulated many point mutations and small deletions, and others retained a high degree of similarity to the unmutated parental haplotype. In this assembled representative sample (#6), a ∼12kb region exhibits a high number of SNVs relative to the surrounding regions.

It has been previously observed that mutations arising from gamma irradiation are not perfectly random (Hase et al., 2020; Li et al., 2022). These patterns may emerge from chromosome conformation, chromatin presence, the effective recruitment of DNA repair proteins, and sequence-specific susceptibility to mutations (Rogozin and Pavlov, 2003; Cowell et al., 2007; Falk et al., 2008; Danforth et al., 2022). Identified novel structural variants (50bp-200kb) were detected in all samples, although these structural variants were present at a much lower rate than small variants (Figure 3b). Some of the identified structural variants were highly complex, with nested variation (Figure 5). Even one structural variant can have significant ramifications for an organism through the insertion, deletion, duplication, or inversion of a gene or regulatory element, altering gene expression, gene product identity, or gene dosage in a way that generates a novel phenotype (Weischenfeldt et al., 2013; Scott et al., 2021). The dense presence of structural variations in all samples means all progeny are likely to have radically altered regulatory pathways, and many genes are likely to be modified or removed altogether. While gene-poor regions of the genome appear to be rich in novel variants, these are far from the only novel variant-rich regions, many of which fall in gene-rich areas. Even the long arm of Chromosome 13, the area with the lowest rate of observed novel structural variants, has many observed novel small variants (Figure 1).

**Figure 5.**
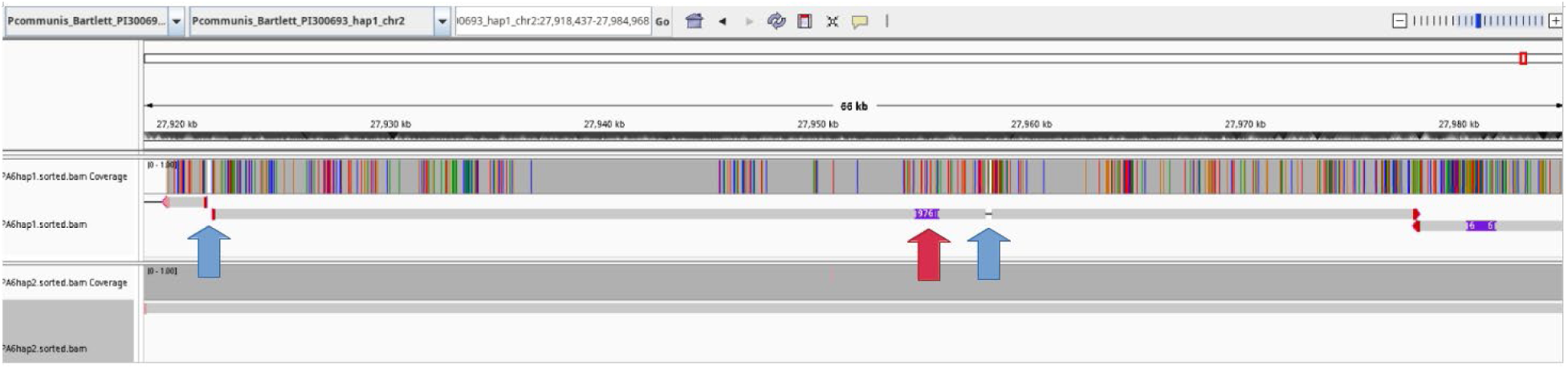
An IGV view of a highly complex nested structural variant in sample #6. A large inversion (∼56,000bp with respect to the reference sequence) on Chromosome 2 contains two major deletions (99bp and 150bp, indicated by blue arrows), two major insertions (68 and 976bp, both indicated by the red arrow), and many SNVs (shown on the coverage track). Arrows at the end of alignment sections indicate directionality of alignments, with this segment reversed relative to the flanking sequence. Reads corresponding to the unmutated haplotype on the bottom track show no consistent variation.

Copy number variations (>200kb deletions or duplications) were considerably less common than smaller variants, but about half of the samples had at least one confidently called large deletion (Table 2). The data did not identify any genomic duplications. Gamma irradiation is known to generate small, rather than large variants preferentially (Li et al., 2019a; Hase et al., 2020). It appears that even at the very high dose of radiation delivered to the pollen in this experiment, this remains true, with only 42 novel large deletions present. To validate at least one of these large deletions, a high coverage sample (PA6 ‘Bartlett’ × ‘Bartlett’(i), 13-3) was assembled and aligned to the reference; the loss of one haplotype in this region was supported (Figure 6). The identified breakages are located near the known centromere positions of the chromosomes they fall on (Sun et al., 2025), which are notably gene-poor (Figure 1).

**Figure 6.**
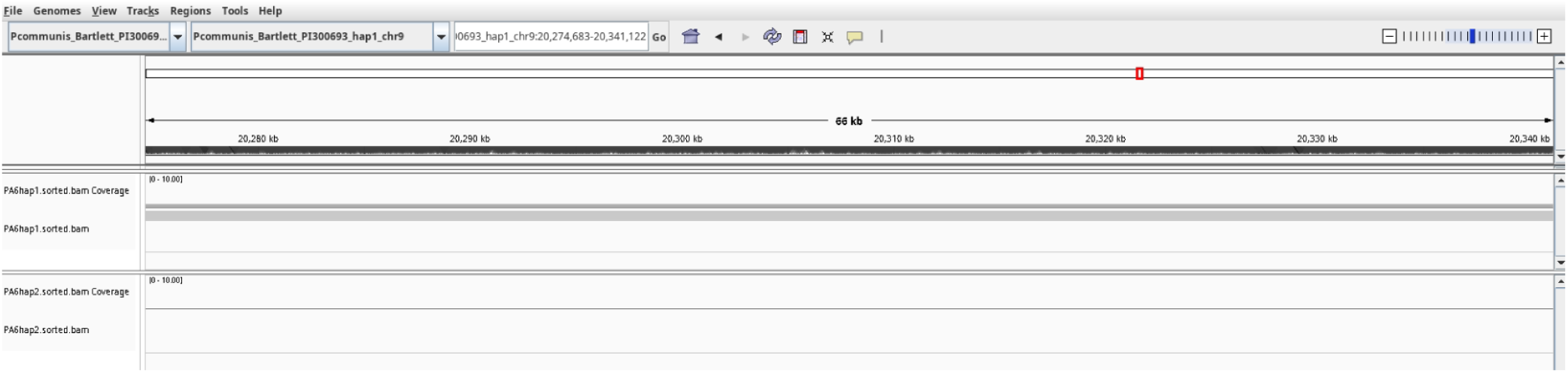
An IGV view of a location with a predicted copy number variation-scale deletion. Presence of only one assembled haplotype observed in the predicted large deletion on Chromosome 9 in Sample #6, supporting the determination of a copy number variant (in this case, a 1.2 Mb deletion). The top track corresponds to a haplotype with no deviation from the reference, while the bottom track is absent.

**Table 2.** Table of copy number variations. Chromosomal locations of large (>200kb) deletions within sequenced samples binned at 25kb resolution.

| Sample | Chromosome | Start Site (Mb) | End Site (Mb) | Length (Mb) |
| --- | --- | --- | --- | --- |
| PA8 | 1 | 21.6 | 23.75 | 2.15 |
| PA22 | 2 | 9.78 | 12.13 | 2.35 |
| PA37 | 3 | 12.3 | 14.48 | 2.17 |
| PA21 | 3 | 12.53 | 15.7 | 3.17 |
| PA32 | 3 | 12.53 | 15.63 | 3.1 |
| PA7 | 4 | 10.45 | 11.73 | 1.27 |
| PA5 | 5 | 23.73 | 26.7 | 2.97 |
| PA8 | 5 | 23.78 | 26.68 | 2.9 |
| PA19 | 6 | 15.45 | 16.55 | 1.1 |
| PA37 | 6 | 16.05 | 18.03 | 1.97 |
| PA8 | 6 | 16.6 | 18.38 | 1.77 |
| PA21 | 7 | 11.23 | 15.78 | 4.55 |
| PA23 | 7 | 11.45 | 15.43 | 3.97 |
| PA5 | 7 | 11.75 | 14.6 | 2.85 |
| PA24 | 7 | 11.9 | 15.43 | 3.52 |
| PA19 | 7 | 12.05 | 15.5 | 3.45 |
| PA30 | 8 | 9.35 | 10.75 | 1.4 |
| PA5 | 9 | 18.18 | 21.43 | 3.25 |
| PA17 | 9 | 18.63 | 20.15 | 1.52 |
| PA19 | 9 | 18.63 | 19.58 | 0.95 |
| PA12 | 9 | 18.7 | 21.3 | 2.6 |
| PA18 | 9 | 18.7 | 21.48 | 2.77 |
| PA32 | 9 | 18.75 | 21.3 | 2.55 |
| PA21 | 9 | 19.05 | 21.35 | 2.3 |
| PA6 | 9 | 20.25 | 21.48 | 1.22 |
| PA32 | 10 | 9.73 | 13.03 | 3.3 |
| PA22 | 11 | 17.2 | 19.73 | 2.52 |
| PA7 | 11 | 18.33 | 20.43 | 2.1 |
| PA32 | 13 | 5.25 | 7.58 | 2.32 |
| PA37 | 13 | 5.25 | 7.53 | 2.27 |
| PA21 | 14 | 13.43 | 16.28 | 2.85 |
| PA3 | 14 | 13.5 | 16.6 | 3.1 |
| PA2 | 14 | 13.85 | 16.08 | 2.22 |
| PA13 | 14 | 13.88 | 16 | 2.12 |
| PA12 | 14 | 14.43 | 15.98 | 1.55 |
| PA17 | 15 | 10.75 | 12.33 | 1.57 |
| PA18 | 15 | 10.75 | 13.38 | 2.62 |
| PA21 | 15 | 10.75 | 14.03 | 3.27 |
| PA23 | 15 | 10.75 | 14.45 | 3.7 |
| PA1 | 15 | 11.28 | 13.1 | 1.82 |
| PA32 | 17 | 15.83 | 17.6 | 1.77 |
| PA37 | 17 | 15.83 | 16.83 | 1 |

Centromeric and pericentromeric regions appear to be hotspots for breakage due to their highly repetitive nature. Pericentromeric regions have been identified as areas of elevated structural variant formation in plants (Du et al., 2012; Bolon et al., 2014). As these regions tend to be gene-poor, it may not immediately impair survival as significantly as large deletions elsewhere in the genes, where changes in gene dosage can lead to lethal defects (Schuster-Böckler et al., 2010; Naish and Henderson, 2024). While mutations to centromeres may result in impaired chromosome segregation and loss of an entire chromosome, even in cases where the original centromere is damaged, proper kinetochore formation and chromosome stability is possible (Nasuda et al., 2005; Xue et al., 2022). *Pyrus communis* is typically diploid and may not tolerate large deletions or the loss of entire chromosomes as readily as many polyploid species (Fox et al., 2020; Vande Zande et al., 2023; Fiala et al., 2024; Morris et al., 2025). *Pyrus* and Malinae, more generally, underwent a whole-genome duplication event in the Paleogene era (Q. Li et al., 2019; Sun et al., 2023; Wu et al., 2013), which may suggest that these duplicated regions are more tolerant of large deletions or other structural variants. Genes duplicated in the whole genome duplication event are well conserved, however, including in terms of gene functionality, making dosage balance still critical for physiological integrity (Li et al., 2019b). Large copy number variants in these regions would likely be lethal. For all variant sizes, there is an apparent inverse relationship between gene density and variant density (Figure 1), with a correlation coefficient of −0.112 for small variants (p<0.001), −0.213 for structural variants (p<0.001), and −0.303 for copy number variants (p<0.001). The pattern of copy number variations observed in this set likely reflects a survivorship bias, with large deletions outside of the observed regions being poorly tolerated.

### Ploidy Analysis

Altered ploidy levels are a common outcome of gamma irradiation. Four samples (#7, #8, #22, #37) display evidence of alternate ploidy levels, three triploids (#7, #8, #22) and one tetraploid (#37). For these samples, the nQuire Gaussian mixture model on read-mapped allele frequencies shows the best fit (lowest value) for a non-diploid model (Figure 7). Higher degrees of ploidy may result from double-strand DNA breaks and other genotoxic stressors inducing endoreduplication, which may have occurred early in embryo development in this sample (Adachi et al., 2011; De Veylder et al., 2011). In mulberry (*Morus alba*), the formation of tetraploids from gamma irradiation was a common result (Katagiri, 1976). Tetraploidy may be advantageous as a precursor for the creation of triploid progeny, which often exhibit enhanced agronomic traits, but also intrinsically for traits such as drought tolerance in rootstock, as seen in tetraploid *Citrus* (Allario et al., 2013; Sedysheva and Gorbacheva, 2013; Howard et al., 2023). Further validation of this alternate ploidy may be warranted by utilizing flow cytometry or karyotyping analysis.

**Figure 7.**
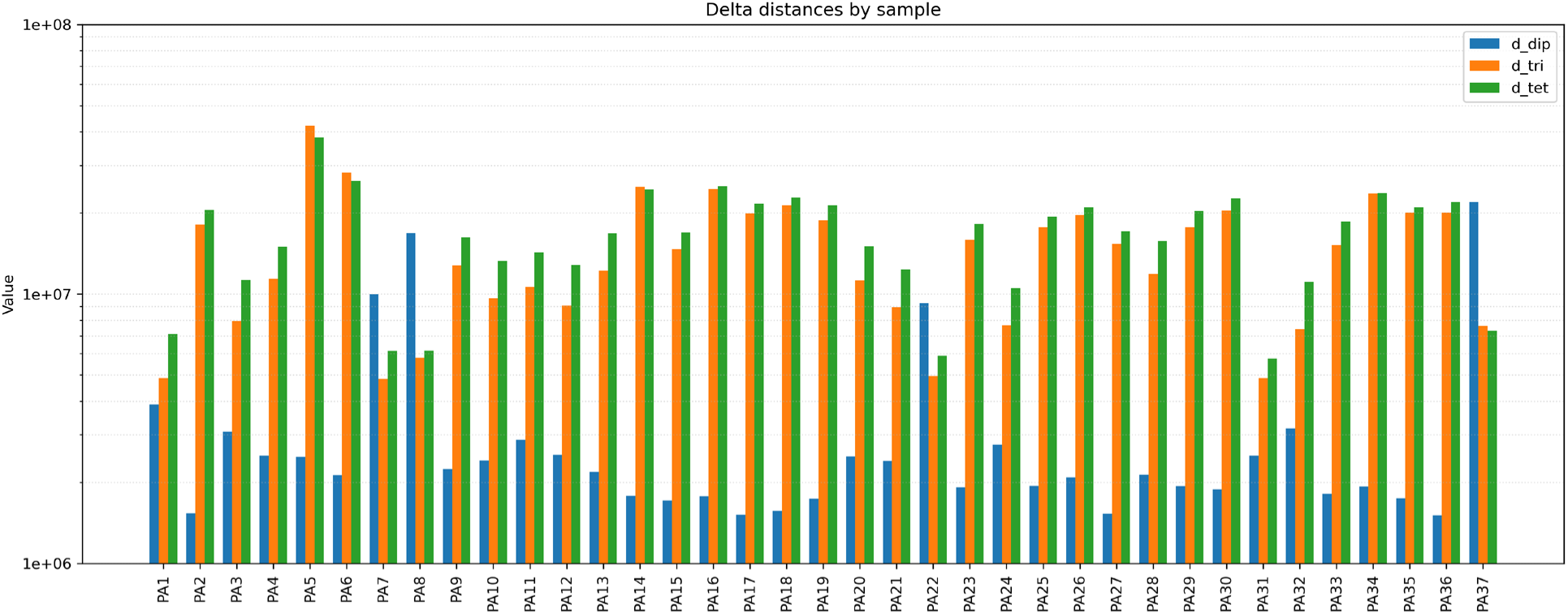
Analysis of ploidy level of all samples. Model fit delta for each sample relative to a diploid (blue), triploid (yellow), and tetraploid (green) model as assessed by nQuire. Model with the lowest delta (best fit) is presumed to describe the ploidy of the sample.

### Mutation Rate

The dose of ionizing radiation delivered to the pollen in this study was 1890 Gy, and the median rate of mutation was 153 small variants/Gy and 0.228 structural variants/Gy in an approximately ∼ 500Mb genome (excluding samples #5 and 31 for reasons discussed below). Owing to the dense pattern of resulting variation, only a small proportion of the seeds yielded were viable, with 49 seeds germinating, and 37 resulting lines surviving at least 10 years. The observed rate of mutation is higher than that seen in gamma-irradiation of *Arabidopsis* dry seed (Kitamura et al., 2022). Vast differences are to be expected due to varying genetic backgrounds, tissue types, radiation doses, and radiation sources. Radiation dose effects are non-linear, especially at high doses, which may also contribute to the observed results (UN FAO/IAEA, 2011; Cornforth et al., 2017; McMahon, 2019; Hase et al., 2020). At nearly 2000 Gy, the DNA repair mechanisms of the target pollen were likely saturated, leading to a non-linear increase in mutations per unit of radiation absorbed (Sánchez-Reyes, 1992; McMahon, 2019; McMahon and Prise, 2019). Pollen appears relatively robust to irradiation compared to vegetative tissues, with high expression of DNA repair genes (Borges et al., 2008; Hirano et al., 2013). However, seeds appear to be harder (Hase et al., 2023). Pollen is capable of germination even at very high radiation dosages over 1000 Gy, with a reported LC50 of 2200 Gy for apple and pear pollen, although embryo viability tends to decrease sharply at doses of >125 Gy (Visser and Oost, 1981; Borges et al., 2008; Hirano et al., 2013). The successful production of seeds, but a very low seed germination rate in this study, supports these observations.

### S-locus Analysis

A total of 15 samples in this study were purportedly derived from self-crosses (Table 1). Gamma irradiation can cause the breakdown of *S*-locus mediated self-incompatibility observed in *Pyrus* and *Malus*, a physiological trait that impedes both the breeding and production of pear fruit (Claessen et al., 2019; Abe et al., 2024; Nishio et al., 2024). Upon inspection of alleles at the SFBB gene at the *S*-locus, however, none of the individuals appeared to be a likely genuine self-cross, containing alleles consistent with only a single parent (Table 3). While some ambiguity exists as to the exact crosses which were made in some instances, as ‘Bartlett’ and ‘d’Anjou’ share the S-101 genotype, and ‘Comice’ and ‘Abbe Fetel’ share both the S-104/S-105 genotype (Takasaki et al., 2006; Goldway et al., 2009; Sanzol, 2009). The reported self-crosses were likely the result of mixed pollen samples from multiple genetic backgrounds, rather than real success in self-pollination. Two individuals had an *S*-locus genotype inconsistent with any of the parental accessions (Lines 5 and 31), raising the possibility that they were pollinated with pollen not part of the experiment at all, which likely explains their large number of genetic variants not seen in any of the parental accessions (Figure 2). As these samples are likely to contain non-parental genetic material, the presence of novel genetic variants may not be adequately assessed by comparison to parental genotypes and have been removed from estimates of overall mutation rate.

**Table 3.** Purported self-crosses (column 2) and the actual crosses the individuals likely arose from (column 4), as assessed by *S*-locus analysis. Identifiers and sample numbers shared with Table 1.

| Sample | Cross | Identifier | Likely Cross |
| --- | --- | --- | --- |
| 4 | 'Bartlett' x 'Bartlett'(i) | 13-1 | 'Bartlett' x 'd'Anjou'(i) |
| 5 | 'Bartlett' x 'Bartlett'(i) | 13-2 | One S-allele not represented by any of the parental accessions |
| 6 | 'Bartlett' x 'Bartlett'(i) | 13-3 | 'Bartlett' x 'Comice'(i) or 'Bartlett' x 'Abbe Fetel'(i) |
| 7 | 'Bartlett' x 'Bartlett'(i) | 13-5 | 'Bartlett' x 'd'Anjou'(i) |
| 8 | 'Bartlett' x 'Bartlett'(i) | 13-6 | 'Bartlett' x 'd'Anjou'(i) |
| 9 | 'Bartlett' x 'Bartlett'(i) | 13-7 | 'Bartlett' x 'Comice'(i) or 'Bartlett' x 'Abbe Fetel'(i) |
| 23 | 'Comice' x 'Comice'(i) | 13-1 | Comice' x 'Bartlett'(i) |
| 24 | 'Comice' x 'Comice'(i) | 13-2 | 'Comice' x 'Bartlett'(i) or 'Comice' x 'd'Anjou'(i) |
| 25 | 'Comice' x 'Comice'(i) | 13-4 | 'Comice' x 'Bartlett'(i) |
| 26 | 'Comice' x 'Comice'(i) | 13-5 | 'Comice' x 'Bartlett'(i) |
| 27 | 'Comice' x 'Comice'(i) | 13-8 | 'Comice' x 'Bartlett'(i) |
| 28 | 'Abbe Fetel' x 'Abbe Fetel'(i) | 13-1 | 'Abbe Fetel' x 'Bartlett'(i) |
| 29 | 'Abbe Fetel' x 'Abbe Fetel'(i) | 13-2 | 'Abbe Fetel' x 'Bartlett'(i) or 'Abbe Fetel' x 'd'Anjou'(i) |
| 30 | 'Abbe Fetel' x 'Abbe Fetel'(i) | 13-3 | 'Abbe Fetel' x 'Bartlett'(i) |
| 32 | 'd'Anjou' x 'd'Anjou'(i) | 2017 | 'd'Anjou' x 'Bartlett'(i) |

### Physiological Outcomes

Reflecting the high degree of mutation found in all samples, none of the surviving lines has flowered yet, despite having lived for 12 years. Genes responsible for the flowering and the initiation of reproductive development are numerous, and single mutations can fail to develop reproductive structures (Weigel et al., 1992; Komiya et al., 2008; Matsubara et al., 2008; Distelfeld and Dubcovsky, 2010; Peng et al., 2015). Consequently, sterility is a common effect of irradiation of plant tissues; in *Arabidopsis*, progeny plant sterility approached 100% as the gamma irradiation dose delivered to pollen approached 1000 Gy (Yang et al., 2004). While high doses such as the 1890 Gy delivered to pollen in this study cause very high rates of sterility (100% in the case of the study), even doses as low as 100 Gy can render the majority of progeny sterile, depending on the stage of pollen maturity (Yang et al., 2004).

Due to the large number of mutations spread across the genome, exact genetic variants that contribute to this lack of floral development cannot be predicted. Regulatory elements can influence genes tens to hundreds of kilobases away in plants, and distant loci seemingly unrelated to specific phenotypes can exert unexpected impacts (Boyle et al., 2017; Mathieson, 2021; Fan et al., 2022; Wang et al., 2024), and on the paternal haplotype, there are regions without extensive mutation. There is no large window free of novel variants in this dataset (Figure 1); this degree of novel mutation makes it effectively impossible to disentangle any predicted gene-phenotype relationships. While these individuals may not be suitable as scion cultivars due to their inability to flower and thus create fruit, there is potential value in testing them as rootstocks. As the exact mechanism of vigor control or induced dwarfing in vigor-controlling rootstocks remains inadequately understood, with disparate proposed mechanisms including carbohydrate partitioning, phytohormone signaling, and partial incompatibility (Basile and DeJong, 2018), testing individuals created through non-directed mutagenesis may be worthwhile.

## Conclusion

Successful mutagenesis was confirmed in all surviving mutation-bred *Pyrus communis* lines subjected to a high dose of ionizing radiation from a Cobalt-60 source. The high degree of novel variation detected in these individuals encompassed variants at all scales, ranging from base substitutions to deletions exceeding 1 Mb. The approximate rate of detected mutations was 153 small (< 50 bp) variants and 0.228 large (> 50 bp) variants per gray of radiation delivered to the pollen at a dose of 1890 Gy. One potential tetraploid was identified. These progenies have not flowered after 12 years, indicating that these individuals are unlikely to contribute to the discovery of functional variation for scion cultivar improvement. However, they may still be suitable as rootstocks or useful for characterizing structural traits. The approach of using a targeted pangenome in the context of tree fruit breeding may be applicable elsewhere as a solution that captures the benefits of pangenomes without encountering the scaling issues that plague extensive pangenomic analyses.

## Author Contributions

June Labbancz: Methodology, Investigation, Formal Analysis, Writing - Original Draft. Nathan Tarlyn: Methodology, Investigation, Writing – Review and Editing. Kate Evans: Conceptualization, Funding acquisition, Writing-Review & editing. Amit Dhingra: Conceptualization, Resources, Supervision, Funding acquisition, Project Administration, Writing - Review and Editing.

## Acknowledgements

The authors thank Mr. Marco Galli, M.S. and Danielle Guizman, M.S. for assistance in performing the crosses, and Daniel Velasquez in the maintenance of seedlings in the greenhouse.

